# Wnt signalling drives context-dependent differentiation or proliferation in neuroblastoma

**DOI:** 10.1101/236745

**Authors:** Marianna Szemes, Alexander Greenhough, Zsombor Melegh, Sally Malik, Aysen Yuksel, Daniel Catchpoole, Kelli Gallacher, Madhu Kollareddy, Ji Hyun Park, Karim Malik

## Abstract

Neuroblastoma is one of the commonest and deadliest solid tumours of childhood, and is thought to result from disrupted differentiation of the developing sympathoadrenergic lineage of the neural crest. Neuroblastoma exhibits intra-and intertumoural heterogeneity, with high risk tumours characterised by poor differentiation, which can be attributable to MYCN-mediated repression of genes involved in neuronal differentiation. MYCN is known to co-operate with oncogenic signalling pathways such as Alk, Akt and MEK/ERK signalling, and, together with c-MYC has been shown to be activated by Wnt signalling in various tissues. However, our previous work demonstrated that Wnt3a/Rspo2 treatment of some neuroblastoma cell lines can, paradoxically, decrease c-MYC and MYCN proteins. This prompted us to define the neuroblastoma-specific Wnt3a/Rspo2-driven transcriptome using RNA sequencing, and characterise the accompanying changes in cell biology.

Here we report the identification of ninety Wnt target genes, and show that Wnt signalling is upstream of numerous transcription factors and signalling pathways in neuroblastoma. Using live-cell imaging, we show that Wnt signalling can drive differentiation of SK-N-BE(2)-C and SH-SY5Y cell-lines, but, conversely, proliferation of SK-N-AS cells. We show that cell-lines that differentiate show induction of pro-differentiation BMP4 and EPAS1 proteins, which is not apparent in the SK-N-AS cells. In contrast, SK-N-AS cells show increased CCND1, phosphorylated RB and E2F1 in response to Wnt3a/Rspo2, consistent with their proliferative response, and these proteins are not increased in differentiating lines. By meta-analysis of the expression of our 90 genes in primary tumour gene expression databases, we demonstrate discrete expression patterns of our Wnt genes in patient cohorts with different prognosis. Furthermore our analysis reveals interconnectivity within subsets of our Wnt genes, with one subset comprised of novel putative drivers of neuronal differentiation repressed by MYCN. Assessment of β-catenin immunohistochemistry shows high levels of β-catenin in tumours with better differentiation, further supporting a role for canonical Wnt signalling in neuroblastoma differentiation.

## Introduction

Neuroblastoma is responsible for approximately 15% of paediatric cancer deaths, with about 40% of patients considered to be high-risk cases with very poor prognosis [1, 2]. Neuroblastoma is a biologically and clinically heterogeneous cancer arising from the sympathoadrenergic lineage of the neural crest [3]. Very recently, studies have demonstrated that tumours and cell-lines derived from them exhibit cellular heterogeneity based on enhancer usage and core regulatory transcriptional circuits [4, 5]. The paradigmatic mechanism for disrupted differentiation in neuroblastoma is contingent on amplification of the *MYCN* proto-oncogene [6], with high levels of MYCN protein leading to direct repression of genes necessary for terminal differentiation in the sympathetic nervous system [7, 8]. As well as *MYCN* amplification (MNA), high risk neuroblastomas have also been shown to elevate telomerase reverse transcriptase (*TERT*) expression through deregulatory genomic rearrangements [9, 10]. Kinase pathways are also deregulated in neuroblastoma, including activating mutations of ALK [11], increased Akt signalling in stage 3 and 4 neuroblastoma [12], and relapsing neuroblastomas displaying mutations in the Ras-MAPK pathway [13, 14].

Another oncogenic pathway reported to be involved in neuroblastoma, as well as many other cancers is canonical Wnt signalling. Here signalling is mediated by secreted Wnt ligands that are bound by membrane receptor complexes including Frizzled proteins and LRP5/6. This then triggers inactivation of a cytoplasmic “destruction complex” that limits cytoplasmic β-catenin, permitting its stabilization and nuclear availability. The TCF/Lef family of transcription factors are then able to utilise β-catenin as a transcriptional co-activator, and instigate target gene expression [15]. Amplification of canonical Wnt signalling can be achieved through the participation of another set of receptors, the leucine-rich repeat-containing G-protein coupled receptors (LGR4, 5, 6) and their ligands, the R-Spondins (Rspos) [16]. LGR-Rspo complexes at the cell membrane decrease the endocytic turnover of Frizzled-LRP5/6 by neutralising the ubiquitin ligases RNF43 and ZNRF3 [17]. Oncogenic Wnt/β-catenin signalling is best exemplified in colorectal cancers where activating mutations in β-catenin or loss of APC (Adenomatosis Polyposis Coli) function (a key destruction complex protein) results in high cytoplasmic and/or nuclear β-catenin [18], constitutive β-catenin-TCF/Lef activity and expression of canonical Wnt target genes, such as *MYC* and *CCND1* [15]. In other cancers such as medulloblastoma, however, Wnt pathway activation is associated with a more favourable clinical subtype distinct from the more aggressive c-MYC driven subtype [19].

In neuroblastoma, mutations in Wnt pathway components have only been reported very recently [20], and identified mutations predicted to have high functional impact in a Wnt geneset defined by the KEGG database, and included *NFATC1*, *FBXW11*, *TP53*, *AXIN2*, *LRP5*, *CCND1*, *FZD9*, *DVL2*, *FOSL1*, *WNT7B*, *VANGL1*, *LEF1*, and *PPP3CB* genes. Interestingly, the latter three gene mutations introduce premature termination, suggestive of a tumour suppressive role of Wnt signalling in neuroblastoma. Other studies in neuroblastoma have suggested that oncogenic deregulation of Wnt signalling occurs, primarily based on over-expression of canonical Wnt pathway target genes identified in other tissues and cancers. Examples include high *FZD1* expression associated with chemoresistance [21], FZD6 marking highly tumorigenic stem-like cells in mouse and human neuroblastoma [22], and FZD2-dependent proliferation of neuroblastoma lines [23]. Furthermore, deregulated Wnt has been suggested to drive the over-expression of *MYC* in non-*MYCN* amplified (non-MNA) high-risk neuroblastomas [24]. Conversely, however, another study utilising chemical agonists and inhibitors of the Wnt pathway has suggested that Wnt signalling hyperactivation directs neuroblastoma cells to undergo apoptosis, and inhibition of Wnt signalling blocks proliferation and promotes neuroblastoma differentiation [25].

Our previous work reported high expression of the Wnt modulator LGR5 in a subset of neuroblastoma cell-lines as well as poorly differentiated primary neuroblastomas [26]. Using a TCF/Lef reporter assay (TOPFLASH), we showed that three LGR5-expressing neuroblastoma cell-lines with different oncogenic drivers, SK-N-BE(2)-C (MNA), SH-SY5Y (*ALK* mutant) and SK-N-AS (*NRAS* mutant) displayed highly inducible β-catenin-TCF/Lef-regulated transcription when treated with recombinant Wnt3a and R-Spondin 2 (Rspo2), with a strong requirement for LGR5/Rspo2 apparent for maximal induction. Although these neuroblastoma cell lines underwent apoptosis after short-interfering RNA (siRNA)-mediated LGR5 knockdown, depletion of β-catenin did not affect cell survival. This suggested that apoptosis after LGR5 depletion occurred independently of Wnt/β-catenin signalling, and further analyses demonstrated a novel pro-survival regulatory influence of LGR5 on MEK/ERK signalling, independent of Wnt/β-catenin signalling [26]. This dual regulatory capacity of LGRs was subsequently also demonstrated in skin carcinogenesis [27].

Although our previous study showed that several established target genes of canonical Wnt signalling were induced in the neuroblastoma cell lines treated with Wnt3a/Rspo2, including *AXIN2* and *LEF1*, we found that *SOX2*, *MYC* and *MYCN*, which were shown to be Wnt pathway target genes studies in other tissue systems (http://web.stanford.edu/group/nusselab/cgi-bin/wnt/), did not exhibit strong induction. In fact, MYC and MYCN protein levels, as well as the activated kinases Akt and ERK, were reduced after Wnt3a/Rspo2 treatment [26]. This, together with the seemingly discrepant literature on Wnt-mediated effects on neuroblastoma cells, prompted us to identify *bona fide* Wnt target genes in neuroblastoma using RNA sequencing of SK-N-BE(2)-C cells treated with Wnt3a/Rspo2, and thereafter correlate the neuroblastoma Wnt signature with clinical parameters. These analyses, together with our evaluation of Wnt3a/Rspo2 effects on neuroblastoma cell biology, reveal that Wnt regulates recently discovered drivers of differentiation such as *EPAS1* [28] and *BMP4* [29] in cell lines that undergo differentiation. Our study also demonstrates further complex interactions between Wnt signalling and other regulatory pathways likely to be involved in neuroblastoma development.

## Materials and Methods

### Neuroblastoma cell lines and culture conditions

Neuroblastoma cell lines were from Carmel McConville (University of Birmingham). The cell lines were grown in DMEM/Nutrient Mixture F-12 Ham (Sigma) supplemented with 10% (v/v) fetal bovine serum (FBS) (Life Technologies), 200 mM L-Glutamine (Sigma), 100 units/ml penicillin, 0.1 mg/mL streptomycin (Sigma) and 1% (v/v) non-essential amino acids (Life Technologies). Prior to stimulation with growth factors, cells were grown in the same medium under low serum conditions, containing 1% (v/v) FBS for 24 hours.

Transient knockdowns were performed by using short interfering RNAs. Briefly, 3 x 10^5^ cells were seeded in 6-well plates after reverse transfection with 50 nM siRNA and RNAiMAX (Invitrogen), both diluted in OptiMEM media (Invitrogen). Sequences for siRNAs are shown in Supplementary table 1.

### Live cell imaging and migration assays

To monitor proliferation in real time, neuroblastoma cells were seeded in a 96-well plate, in quadruplicates, in media containing 1% (v/v) FBS (5000 cells/well). After incubation for 24 hours, cells were treated with recombinant Wnt3a and/or R-Spondin2 (R&D Systems, 100 ng/mL each) using low serum conditions and transferred to the IncuCyte ZOOM live cell imaging system (Essen BioScience). Epidermal growth factor (Sigma) was used at 10 ng/mL. Images were taken in four different fields in each well, every two hours and phase confluence was calculated as a surrogate for growth at each time point. To measure cell death, DRAQ7 (BD Biosciences), a dye that is taken up by dead cells only and fluoresces at 678 nm, was added at 1.5 μM concentration and fluorescence was measured in the red channel. To measure migration, cells were seeded at near confluent levels (50000 cells/well) in ImageLock 96-well plates (Essen BioScience), in quadruplicates, in media containing 1 % FBS (v/v). Cells were stimulated with Wnt3a and/or R-spondin2 24 hours later, and, following another 24 hours, scratched by using the WoundMaker tool (Essen BioScience) according to manufacturer’s instructions. Migration was monitored at every 2 hours automatically, and wound width and relative wound density were calculated for every time point.

### RNA sequencing and bioinformatic analyses

Serum-starved SK-N-BE(2)-C cells were stimulated with 100 ng/mL Wnt3a and R-spondin2 in duplicates, for 6 hours and were subsequently harvested. Control cells were treated with the vehicle, 0.2 % (w/v) BSA in PBS. RNA was extracted by using miRNeasy Mini Kit (Qiagen), according to manufacturer’s instructions. RNA concentration and quality was checked by using Nanodrop spectrophotometer and Bioanalyzer (Agilent). All RNA integrity values (RIN) were above 9. cDNA libraries were prepared from 1 ug RNA as template, using the TruSeq Stranded Total RNA Library Prep Kit (Illumina) according to manufacturer’s instructions. The libraries were sequenced by using the paired-end option with 100 bp reads on Illumina HiSeq 2000 and min. 50 million reads were obtained per sample. Formatting and quality control (FASTQC v0.11.5, Babraham Institute) were performed by using the Galaxy platform [30] installed at the University of Bristol. The paired end reads were then aligned to the human genome (hg38) by using TopHat2 (v2.0.14) [31]. The resulting alignment files (BAM) files were uploaded to SeqMonk v1.38 (https://www.bioinformatics.babraham.ac.uk/projects/seqmonk/). Gene expression was quantified by using the Seqmonk RNA-seq analysis pipeline, with the option of choosing reads along mRNA features as a basis of quantification. Differentially expressed genes (DEG) were identified as showing significant changes by either DESEQ2 [32] or Intensity Difference Testing (Seqmonk) algorithms (p<0.005) between Wnt3A/Rspo2 treated and control sample sets, and having a fold difference of minimum 1.3. RNA sequencing data is available from the European Nucleotide Archive (ENA) under the study accession number PRJEB21488.

We performed Gene Signature Enrichment Analysis (GSEA) [33] on a preranked list of relative gene expression values (log2-transformed average Wnt3A/Rspo2 treated per control) with the classic weighting option, as recommended for RNA-seq data (Broad Institute). Analysis of gene set interconnectivity and enrichment of Gene Ontology (GO) categories was performed by using tools of the STRING database [34] (http://www.string-db.org). Gene expression and gene set correlations in published RNA-seq and microarray data sets were analysed by using the R2 Genomics Analysis and Visualization Platform (http://r2.amc.nl). Clustering of Wnt target gene expression in the SEQC neuroblastoma RNA-seq data set was performed by using k-means clustering in R2. The option of k=4 provided the best separation of clusters. Wnt metagene groups were defined based on the the similarity of co-expression of the Wnt target genes calculated during clustering. Wnt metagenes were defined as having an average expression of the members of the metagene group. Kaplan Meier survival analysis, indicating the prognostic value of the expression of genes or metagenes was performed by using the Kaplan scan tool in R2. Briefly, Kaplan scan selected the expression threshold providing the best statistical significance of the separation of outcomes associated with high and low expression of a gene or metagene, using the log-rank test.

### RNA extraction and qRT-PCR

RNA was extracted using the miRNeasy Mini Kit (Qiagen) according to manufacturer’s instruction and subsequently transcribed into cDNA by using the Thermoscript kit (Invitrogen). Quantitative PCR was performed on an Mx4000P machine (Stratagene) using QuantiNova kit (Qiagen) and analysed according to the comparative Ct method, normalizing values to the housekeeping gene TBP. Primer sequences are shown in Supplementary Table 1.

### Immunoblotting and Immunohistochemistry

Immunoblotting was essentially as described previously [35]. The antibodies used were the following: E2F1 and MYCN (Santa Cruz, SC-251 and SC-53993 respectively, 1:1000), BMP4 (4680s), CCND1 (2978s), EPAS1 (7096s), NGFR/p75NTR (8238S) phospho-RB (8516s), all Cell Signalling, 1:1000) and beta-actin-peroxidase (A3854, Sigma, 1:40000). Peroxidase conjugated secondary antibodies (Sigma) were diluted 1:5000 and the blots were developed using LumiGLO Peroxidase Chemiluminescent substrate (KPL).

Immunohistochemistry was conducted essentially as described previously [35]. Neuroblastomas were collected at The Children’s Hospital at Westmead Histopathology Department (Sydney, Australia). All human tissues were acquired with appropriate local research ethics committee approval. The β-catenin antibody (Cell Marque, 760-4242) was used after pH8.5 heat retrieval. Staining excluded post-treatment samples which displayed extensive necrotic areas.

## Results

### Wnt3a/Rspo2 drives a unique transcriptional program in neuroblastoma, affecting multiple pathways

We characterised Wnt3a/Rspo2-induced early transcriptomic changes in the MNA neuroblastoma cell line, SK-N-BE(2)-C. Using a bioinformatic cut-off of p=<0.005 along with a minimum fold change of 1.3, RNA-sequencing identified 90 genes showing significant changes in expression (Figure 1A). Eighty-five of the differentially expressed genes (DEGs) were upregulated, with 5 genes showing decreased expression after Wnt3A/Rspo2 treatment (*ASCL1*, *RGS16*, *HS6ST1*, *NME2* and *TMEM108-AS1*).

**Figure 1.**
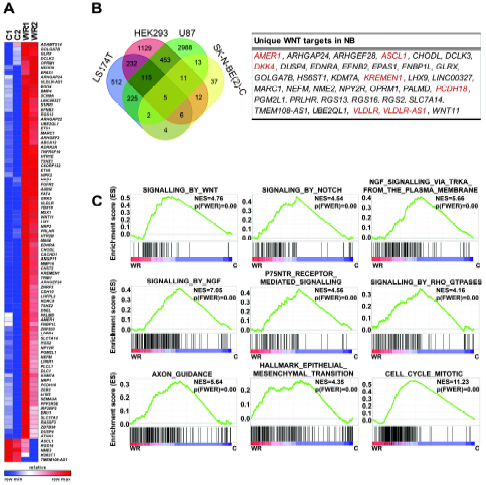
Identification of novel Wnt target genes in neuroblastoma. **(A)** Heatmap depicting 90 differentially expressed genes (DEGs) in Wnt3A/Rspo2 treated SK-N-BE(2)-C cells (WR) at 6 hours vs. control (C), as determined by RNA-seq (n=2). **(B)** Venn diagram comparing the 90 Wnt3A/Rspo2-regulated genes in SK-N-BE(2)-C to Wnt target genes in colorectal cancer (LS174T), glioma (U87) and embryonic kidney cells (HEK293), previously determined by using ChIP-seq. The table lists thirty-seven Wnt target genes uniquely identified in neuroblastoma cells. Wnt feedback/negative regulator genes are shown in red. **(C)** Gene Set Enrichment Analysis (GSEA) of gene expression changes in Wnt3A/Rspo2-induced SK-N-BE(2)-C cells at 6 hours, using the Reactome and Hallmark databases. Respective corrected p values (p FWER) and normalized enrichment scores (NES) are shown. WR, treated; C, control.

In order to assess whether our 90 DEGs had been previously identified as direct targets of β-catenin/TCF-/Lef in other cell types, we compared our gene list with published target genes of canonical Wnt signalling, based on using ChIPseq in U87 glioma cells [36], HEK293, and the colorectal cancer cell line LS174T [37]. Fifty-three out of the 90 DEGs were identified as having β-catenin and/or TCF7L2/TCF4 bound to their promoters in one or more of these three data sets (Figure 1B and Supplementary Table 2). To our knowledge, however, Wnt-dependent increase of transcription has not been demonstrated for many of these 53 genes, including *LIX1*, *S1PR1*, *FAT4*, *HTR1E*, *ETV6*, *TSHZ1* and *NEDD9*. As often seen with induction of Wnt signalling, several genes encoding negative regulators of the pathway are observed amongst the DEGs, including *AXIN2*, *NKD1*, *TNFRSF19*, *ZNRF3*, *KREMEN*, *AMER1* and *VLDLR*. Of the 37 genes not encompassed by the CHiP-seq datasets, *EFNB2* [38], *DKK4* [39] and *WNT11* [40] have been previously shown to be induced by Wnt signalling, and all genes exhibited TCF7L2 binding according to ENCODE/SYDH ChIPseq data (data not shown). Thus our dataset identifies novel Wnt target genes likely to have roles in cell-fate decisions in neuronal development and neuroblastoma.

We found genes encoding neuronal proteins were induced by Wnt3a/Rspo2, including several neurotransmitter receptors (*HTR1E*, *HTR2B*, *NPY2R*, *ADRA2A* and *OPRM1*), neurofilament protein M (*NEFM*), and transcription factors involved in differentiation of the neural crest, including *MSX1* and *MSX2* [41]. *EPAS1*/*HIF-2α*, which was very recently associated with differentiation of neuroblastoma [28] was also upregulated after treatment. *ASCL1*, which is one of the few down-regulated genes amongst the DEGs, is also involved in neuronal differentiation [42], with mRNA downregulation also observed with retinoic acid induced differentiation of neuroblastoma cells [43]. Numerous other transcription factors were also induced such as *ETS1*, *ETV6*, *LHX9*, *TSHZ1*, *TSHZ2*, *ZEB2* and *ZNF503*, indicating complex transcriptional networks triggered by Wnt signalling. Functional annotations for the 90 DEGs are shown in Supplementary Table 2. Analysis of Wnt3a/Rspo2-induced global gene expression changes using Geneset Enrichment Analysis (GSEA) further confirmed induction of neural differentiation pathways (TrkA, nerve growth factor (NGF), NGFR/p75NTR, axon guidance) (Figure 1C). In addition GSEA highlighted influences on numerous signalling pathways, including Wnt, Notch, and Rho GTPase (Figure 1C). Further pathways and processes affected included Hedgehog, mTORC1, and EGFR signalling, as well as glycolysis, oxidative phosphorylation and mRNA splicing (Supplementary Table 3). Epithelial-mesenchymal transition (EMT), which is integral to neural crest differentiation, was also apparent, and, consistent with our previous observation of increased Ki67 following Wnt3a/Rspo2 treatment, effects on the cell cycle and mitosis were evident (Figure 1C) [26].

Using qRT-PCR, we validated 27 of the DEGs in SK-N-BE(2)-C cells and in two other Wnt3a/Rspo2 responsive neuroblastoma cell lines with different driver mutations, SK-N-AS (*NRAS* mutant) and SH-SY5Y cells (*ALK* mutant) following 6 hours and 24 hours of Wnt3a/Rspo2 treatment. While many DEGs were also upregulated (greater than 1.3-fold at 6 hours) in the other two cell lines, the Wnt response in SK-N-AS was more dissimilar and shared 63% of target genes with SK-N-BE(2)-C, compared with 89% for SH-SY5Y (Supplementary Figure 1). Generally, we observed gene induction at 6 and 24 hours, indicating a sustained transcriptional response in SK-N-BE(2)-C and SH-SY5Y. *EPAS1*, *FAT4*, *KREMEN1* were not induced in SK-N-AS cells at either time-point (Supplementary Figure 1). R-spondin 2 alone did not significantly affect the expression of these genes (data not shown).

Taken together, analyses of our novel neuroblastoma Wnt target genes suggests that Wnt signalling may have profound effects on differentiation as well as proliferation via a complex orchestration of downstream transcription factors and signalling pathways.

### Wnt signalling affects growth and differentiation in a context-dependent manner

As our transcriptomic data showed effects on differentiation associated genes such as *ASCL1*, *BMP4*, *NEFM*, and *EPAS1*, and GSEA also indicated stimulation of neuronal differentiation pathway genesets, we next assessed the phenotypic effects of Wnt signalling on SK-N-BE(2)-C, SH-SY5Y and SK-N-AS cell-lines. As shown in Figure 2A, treatment of SK-N-BE(2)-C cells with Wnt3a/Rspo2 induced morphological changes indicative of differentiation, including a more mesenchymal appearance and increased neurite-like projections. Using real-time, live-cell imaging, we quantified changes in cell confluence, and observed no significant proliferative effects with Wnt3a or Wnt3a/Rspo2, in contrast to the positive control EGF (Figure 2B, left panel). However, a significant increase (p=<0.01) of neurite length and neurite branch points was apparent in Wnt3a/Rspo2 treatments, beginning around 24 – 30 hours. Wnt3a alone also induced characteristics of neural differentiation in SK-N-BE(2)-C cells, but at later time points (33 – 39 hours) and at a lesser level (p=<0.05) (Figure 2B, middle and right panels). This is consistent with the observed changes corresponding to Wnt signalling amplitude, which we showed were increased with Wnt3a/Rspo2 co-treatment [26].

**Figure 2.**
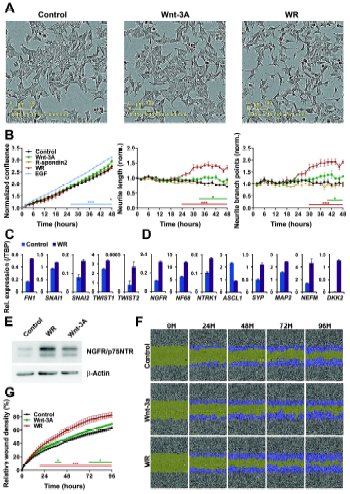
Differentiation and migration of SK-N-BE(2)-C cells after Wnt3A/Rspo2 induction. **(A)** Phase contrast microscopy of control, Wnt and Wnt3A/Rspo2 treated SK-N-BE(2)-C cells at 48 hours. **(B)** Phase confluence was monitored as a surrogate for growth using IncuCyte live-cell imaging. EGF treatment was used as a positive control for proliferation. Neurite lengths and neurite branch points were quantified by using IncuCyte NeuroTrack software. Significance was evaluated at every time point by using T tests at p<0.01 (***) and p<0.05 (*). **(C)** qRT-PCR of transcription factors driving epithelial-to-mesenchymal transition (EMT) in Wnt3A/Rspo2 induced cells at 24 hours. **(D)** qRT-PCR of neuroblastoma differentiation markers in Wnt3A/Rspo2 induced cells at 72 hours. **(E)** Western blot of NGFR/p75NTR in Wnt3A, Wnt3A/Rspo2 and control cells at 72 hours. β-actin was used as loading control. **(F)** Representative fields of a 96-well scratch wound assay, measuring migration after Wnt3A and Wnt3A/Rspo2 treatment vs control (n=4). The original wound is shown in green and the migrating cells are highlighted in blue. **(G)** Relative wound density plot, showing cell confluence within the wound area corrected for proliferation by Incucyte software (n=4). Significance was tested at every time point by using T tests at p<0.01 (***) and p<0.05 (*).

Given the morphological changes observed, and our GSEA analyses, we next evaluated markers of EMT and neural differentiation in Wnt3a/Rspo2 treated cells. Induction of fibronectin (*FN1*), as well as the EMT transcription factors *SNAI1*, *SNAI2*, *TWIST1* and *TWIST2* was apparent in SK-N-BE(2)-C cells 24 hours after Wnt3A/Rspo2 treatment (Figure 3C). Analysis of genes that are increased with retinoic acid induced differentiation of neuroblastoma cells (*NGRF*, *NF68*, *NTRK1*, *SYP*, *MAP2*, *NEFM*, *DKK2*) [44-46] showed that all were elevated after 72 hours Wnt3a/Rspo2 treatment. Downregulation of *ASCL1* was also observed (Figure 3D). The parallels between Wnt and retinoic acid effects were further supported by comparing our DEGs with an array analysis of SH-SY5Y cells, which shows that many Wnt genes are similarly altered by retinoic acid (Supplementary Figure 2). Differentiation was also verified at the protein level for NGFR/p75NTR, which was markedly increased after Wnt3a/Rspo2 treatment, with a moderate increase also observed with Wnt3a alone (Figure 2E). As we observed EMT-like changes, and differentiation induced by all-trans-retinoic acid (ATRA) has also been shown to increase cell migration [47], we evaluated the effects of Wnt induction in a wound-healing/scratch assay. Consistent with our mRNA marker data, Wnt3a/Rspo2 treatment led to increased migration (Figure 3F) and a significantly increased wound cell density from 22 hours post-treatment (p=<0.01, Figure 3G), with more moderate effects also observed with Wnt3a treatment alone at later time points. Together, these results are consistent with Wnt3a/Rspo2 leading to differentiation of SK-N-BE(2)-C cells akin to that induced by retinoic acid.

**Figure 3.**
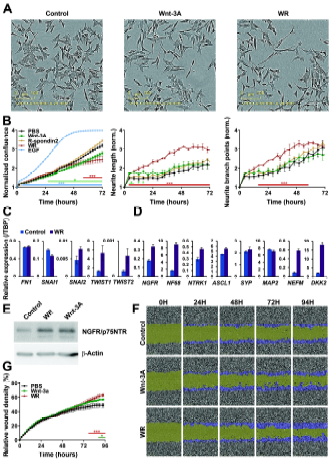
Differentiation and migration of SH-SY5Y cells after Wnt3A/Rspo2 induction. **(A)** Phase contrast microscopy of control, Wnt and Wnt3A/Rspo2 treated SH-SY5Y cells at 48 hours. **(B)** Phase confluence was monitored as a surrogate for growth and EGF was used as a positive control. Neurite lengths and neurite branch points were calculated by using IncuCyte NeuroTrack software at every timepoint and significance was evaluated by using T tests at p<0.01 (***) and p<0.05 (*). **(C)** qRT-PCR of transcription factors driving EMT in Wnt3A/Rspo2 induced cells at 24 hours. **(D)** qRT-PCR of neuroblastoma differentiation markers in Wnt3A/Rspo2 induced cells at 72 hours. **(E)** Western blot of NGFR/p75NTR in Wnt3A, Wnt3A/Rspo2 and control cells at 72 hours. β-actin was used as loading control. **(F)** Representative fields of a 96-well scratch wound assay, measuring migration after Wnt3A and Wnt3A/Rspo2 treatment vs control (n=4). The original wound is shown in green and the migrating cells are highlighted in blue. **(G)** Relative wound density plot, showing cell confluence within the wound area corrected for proliferation by Incucyte software (n=4). Significance was tested at every time point by using T tests at p<0.01 (***) and p<0.05 (*).

We performed similar analyses with SH-SY5Y cells and observed comparable morphological changes to those seen with SK-N-BE(2)-C cells (Figure 3A). However, real-time quantification of cell confluence revealed that, in contrast to SK-N-BE(2)-C cells, the growth of SH-SY5Y cells was inhibited by Wnt3a alone from 26 hours onwards (p=<0.05). Dual treatment with Wnt3a/Rspo2 also led to growth inhibition beginning at about 26 hours and becoming significant at about 56 hours (p=<0.01). Although more delayed, the extent of growth inhibition was greater with the dual treatment (Figure 3B, left panel). Both Wnt3a and Wnt3a/Rspo2 treatments led to an increase in neurite length, but the Wnt3a effect attenuated after approximately 16 hours, whereas the Wnt3a/Rspo2 treated cells continued to show increasing neurite length and also increasing branch points from 16 hours onwards. Of the EMT markers, *SNAI2* and both *TWIST* genes were induced after Wnt3a/Rspo2 treatment (Figure 3C), and induction of neural differentiation genes was also clearly apparent (Figure 3D). In SH-SY5Y cells, similar levels of NGFR/p75NTR induction were observed at the protein level between Wnt3a and Wnt3a/Rspo2 treatments (Figure 3E), and both treatments led to increased cell migration (Figs. 3F, G). Together these data support the differentiation inducing influence of Wnt signalling on SH-SY5Y cells, similar to that observed in SK-N-BE(2)-C cells.

We also evaluated the effects of Wnt3a/Rspo2 treatment on a third LGR5-positive neuroblastoma cell line, SK-N-AS. In marked contrast to SK-N-BE(2)-C and SH-SY5Y cells, we observed no morphological changes after treatments, as well as a robust increase in proliferation (Supplementary Figure 3A, B). Consistent with the lack of visible phenotypic changes, most differentiation markers did not increase, as assessed by qRT-PCR, and some of them decreased. Further, in contrast to SK-N-BE(2)-C and SH-SY5Y cells, NGFR/p75NTR protein negatively correlated with Wnt signalling amplitude, reducing with Wnt3a alone and decreasing further still with Wnt3a/Rspo2 treatment (Supplementary Figure 3C, D). In contrast to a previous report [25], we did not observe Wnt signalling associated cell death (Supplementary Figure 2E). We further explored whether the proliferative response of SK-N-AS cells may be characteristic of their more mesenchymal state, as defined recently [4], by assessing proliferation in another mesenchymal line, GI-MEN. However, Wnt3a/Rspo2 treatment did not trigger proliferation in this line (Supplementary Figure 3F).

To further evaluate the growth and differentiation variance of these cell lines at the protein level, we examined whether induction of two recently characterised inducers of differentiation, BMP4 [29] and EPAS1 [28], corresponded with the propensity of cells to differentiate. As shown in Figure 4A, although BMP4 and EPAS1 were induced by Wnt signalling in the differentiating cell-lines (SK-N-BE(2)-C and SH-SY5Y), their levels in SK-N-AS cells were unchanged. In contrast, assessment of cell-cycle/proliferative markers demonstrated an increase in CCND1 and E2F1 only in SK-N-AS cells. Consistent with our real-time differentiation and proliferation assays, SH-SY5Y cells showed decreased CCND1, E2F1 and phosphorylated RB proteins (Figure 4B).

**Figure 4.**
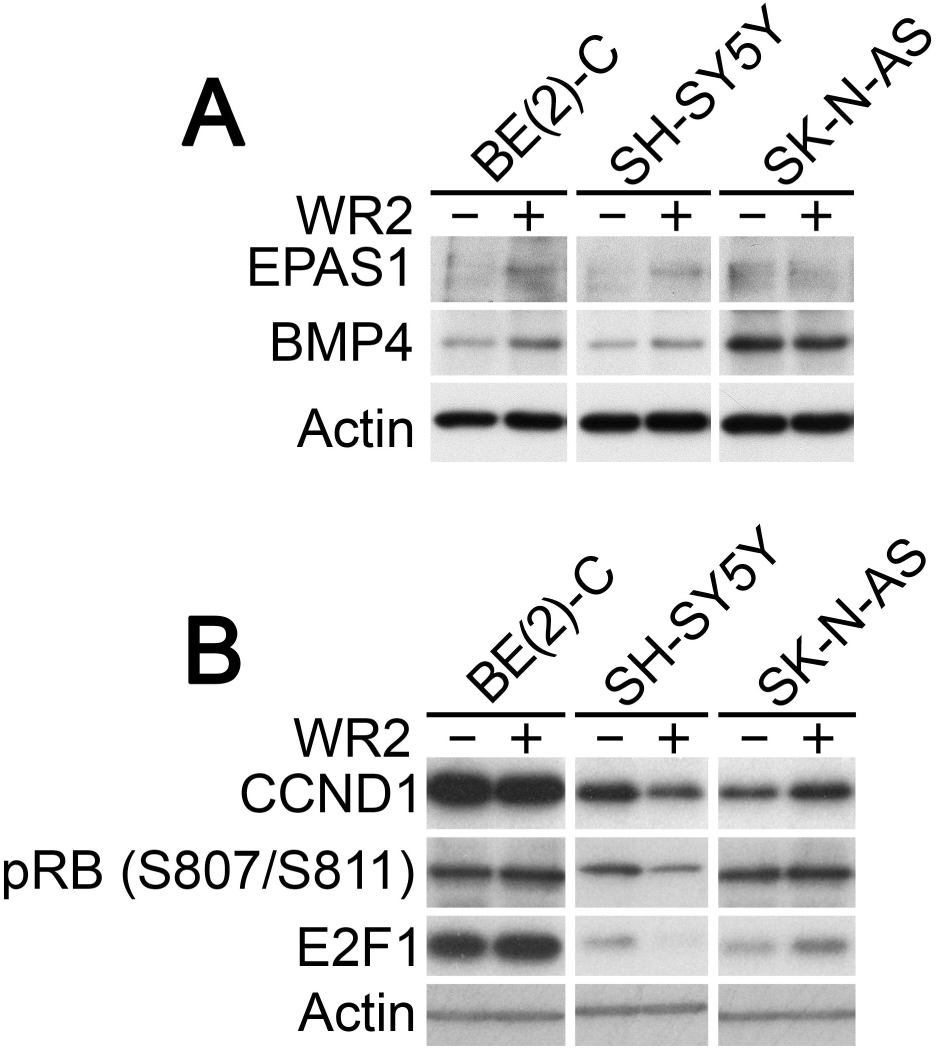
Induction of pro-differentiation and proliferation proteins by Wnt3a/Rspo2 treatment. **(A)** Immunoblotting of BMP4 and EPAS1 after 48 hours of treatment shows increases in SK-N-BE(2)-C and SH-SY5Y cell-lines, but not in SK-N-AS. **(B)** Immunoblotting of CCND1, E2F1 and phospho-RB shows increases in SK-N-AS, but not in SK-N-BE(2)-C and SH-SY5Y cell-lines.

Together our phenotypic analyses show that Wnt signalling can induce a more neuronal phenotype in SK-N-BE(2)-C and SH-SY5Y cells, with no increase in cell proliferation. In SK-N-AS cells, however, quite the opposite was apparent, with Wnt signalling inducing cell proliferation without differentiation.

### Primary neuroblastoma gene expression datamining reveals that Wnt target genes are associated with distinct prognostic groups

Having shown that Wnt induction can prompt differentiation and growth suppression, or in the case of SK-N-AS cells, proliferation, we next evaluated the clinical correlations of our 90Wnt pathway DEGs in publicly available primary neuroblastoma expression data. Firstly, using k-means clustering, we assessed whether patterns of expression of the DEGs existed in a large neuroblastoma cohort (SEQC, n=498, [48]). Strikingly, this analysis revealed that the cohort could be partitioned into 4 clear patient clusters, each with a distinctive expression pattern of our DEGs; in turn, the DEGs could also be grouped into 4 sets, from which we could derive 4 model genes, or Wnt metagenes (WMG-1, −2, −3 and −4) for further analyses (Fig 5A). Such metagenes enable using a single value to represent the combined expression values of multiple co-expressed genes.

**Figure 5.**
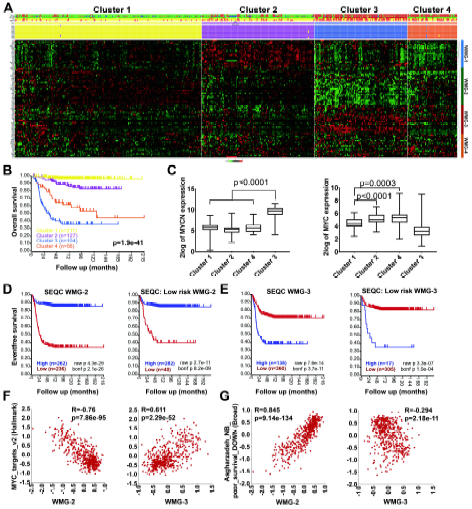
Expression patterns of the 90 identified Wnt target genes in primary neuroblastomas. **(A)** K-means clustering in 498 neuroblastoma tumours (SEQC data set). In the heatmap red indicates high, while green shows low expression. Annotation of patient samples are shown at the top in the following order: risk (high: red, low: green), INSS stage (1: green, 2: dark green, 3: orange, 4: red and 4S: blue), MYCN amplification (MNA: red, non-MNA: green), outcome (survival: yellow, died of disease: red) and progression (no progression: yellow, relapse: red). The result of each iterative round of clustering with k=4 is indicated with rows of yellow, purple, blue and orange coloured squares. Clustering highlighted 4 groups of co-expressed genes that were represented with metagenes (WMG-1 to 4, shown on the right). **(B)** Kaplan Meier analysis (log rank test) of the four patient clusters exhibiting significantly different survival. **(C)** Expression of *MYCN* and *MYC* in the 4 neuroblastoma clusters. Significance assessed by T tests. **(D)** Kaplan Meier survival curves associated with expression of WMG-2 in SEQC data set. All patients and low-risk patients are shown separately. Bonferroni-corrected p values of log rank test are shown. **(E)** Kaplan Meier analysis of WMG-3 as in (D). **(F)** Correlation of WMG-2 and 3 with MYC target genes and **(G)** Asgharzadeh’s set of downregulated genes in non-MNA metastatic neuroblastoma (curated gene sets, Broad Institute) in SEQC. Correlation coefficients (R) and p values are shown.

Of the 4 patient clusters, cluster 3 (blue) and 4 (orange) had high representation of high-risk neuroblastoma (85%, 137/160), as defined by the SEQC dataset, with cluster 3 being composed of mainly MNA tumours (81%, 84/104). This demonstrates the efficacy and veracity of the k-means clustering, which is further supported by the Kaplan-Meier analysis of our k-means derived clusters, where poor overall survival could be ranked as cluster 3>cluster 4>cluster2>cluster 1 (Fig 5B). The expected association of cluster 3 with *MYCN* over-expression was apparent and *MYC* expression was significantly higher in cluster 4, the second cohort associated with very poor prognosis (Fig 5C). We also tested correlations with other neuroblastoma oncogenes, *TERT* and *LIN28B*. These were also higher in clusters 3 and 4 designated by our analyses (Supplementary Figure 4).

With regard to metagene expression patterns between the patient clusters, contrasting patterns of high or low expression were clearly apparent. In the predominantly MNA cluster 3, WMG-2 genes were expressed at low levels compared to other clusters; conversely WMG-3 genes were highly expressed in cluster 3 (Figure 5A). Kaplan-Meier analysis of WMG-2 in four neuroblastoma datasets demonstrated that low WMG-2 expression is an indicator of poor prognosis (Fig 5D, left), Supplementary Figure 5A-C). Notably, although we identified WMG-2 in the predominantly MNA cluster 3, Kaplan-Meier analysis of low-risk tumours in the SEQC dataset (n=322) with WMG-2 demonstrated a highly significant association of low WMG-2 with poor survival (p=8.2 x 10^−9^) (Figure 5D, right). Further assessment of low-risk, non-MNA amplified neuroblastomas across several databases confirmed that a WMG-2(low) signature may help in identification of patients with poor prognosis, despite being classified as lower risk (Supplementary Figure 5A-C).

Conversely, high expression of WMG-3 was associated with poor prognosis across the four datasets, and a WMG-3(high) signature could also identify low risk patients with poor prognosis, albeit with less specificity (Figure 5E, Supplementary Figure 5D-F). The prognostic value of WMG-1 was not as strong as that of WMG-2 or WMG-3, but high expression was associated with better prognosis, after exclusion of MNA tumours. Although this trend was apparent in other datasets, it did not always reach the high levels of significance seen with WMG-2 and WMG-3 (Supplementary Figure 5G). The WMG-4 metagene did not have clear prognostic value, as would be expected from the lack of clear expression pattern observed for the constituent genes (Figure 5A, Supplementary Figure 5H).

As WMG-2 and WMG-3 show distinctive and opposite expression patterns in the predominantly MNA cluster 3, we assessed the correlations of these metagenes with a MYC target gene dataset. In the SEQC data set, a strong inverse correlation (R=-0.76, p=7.86 x 10^−95^) was observed between the Hallmark MYC target genes and WMG-2 (Figure 5F). WMG-3, as expected, had a strong positive correlation with MYC target genes (R=0.61, p=2.29 x 10^−52^), although the two gene sets did not have any genes in common. Remarkably, WMG-2 also had a striking positive correlation (R=0.85, p=9.14 x 10^−134^) with a published gene set representing genes downregulated in poor prognosis non-MNA neuroblastoma (Asgharzadeh_NB_poor_survival_DOWN) [49], despite only sharing one gene (*PGM2L1*) (Figure 5G). An inverse correlation of the Asgharzadeh gene set with WMG-3 was also apparent, but less significant, as expected given that this geneset is limited to non-MNA tumours. Together this reinforces that high expression of the WMG-2 metagene is an indicator of good prognosis regardless of *MYCN* status

### Immunohistochemical analysis of β-catenin in primary neuroblastomas

Our transcriptomic and functional data, together with recent mutation analysis [20], indicate that a canonical Wnt programme may be more active in lower stage neuroblastoma and decreased/suppressed in higher stage neuroblastomas. We therefore examined β-catenin protein expression at the cellular level in neuroblastoma tissue sections. As shown in Figure 6A, poorly differentiated non-MNA neuroblastomas showed very weak or negative staining, and a MNA poorly differentiated tumour exhibited a low level of focal membranous positivity (Figure 6B). In contrast to these stroma poor tumours, stroma rich neoplasms, ganglioneuroblastomas and ganglioneuromas, showed membranous β-catenin staining of differentiating neuroblasts (Figure 6C) and maturing and mature ganglion cells (Figure 6C and 6D), which were embedded in Schwannian stroma. Interestingly, Schwannian stroma displayed intense and extensive cytoplasmic and membranous β-catenin positivity in the Schwann cells, as well as some nuclear staining (Figure 6C and 6D). In these, stroma rich tumours, Schwann cell condensation and increased β-catenin levels were detected around ganglion cells (Figure 6D). Taken together, these results are supportive of our transcriptomic and functional data suggesting that Wnt signalling may prompt differentiation signals for neuroblasts, and further suggest that Schwann cells, which have been shown to promote neuroblast differentiation [50, 51], may be the source of Wnt3a/Rspo2-induced signalling ligands.

**Figure 6.**
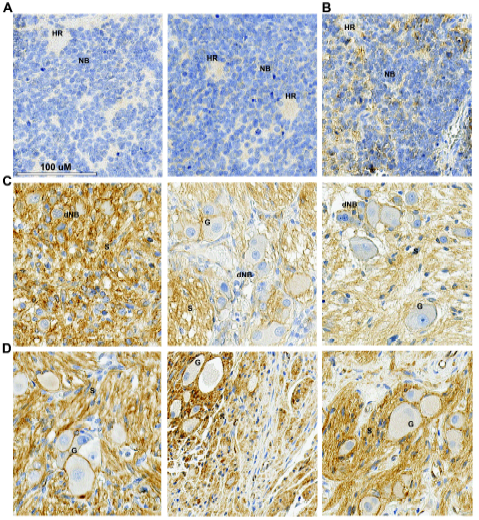
Immunohistochemistry of β-catenin in primary neuroblastomas. **(A)** Poorly differentiated neuroblastoma, non-MNA. **(B)** poorly differentiated neuroblastoma, MNA **(C)** Ganglioneuroblastoma and maturing ganglioneuroma. **(D)** Ganglioneuroma. Labelled are HR, Homer-Wright rosettes; NB, neuroblasts; dNB, differentiating neuroblasts; G, Ganglion cells; S, Schwannian Stroma.

### The influence of MYCN on Wnt signalling and other potential cross-talk

Our transcriptomic clustering (Figure 5A) revealed that certain groups of genes had a very distinctive distribution pattern between the four patient clusters. In particular genes comprising WMG-2 were expressed at markedly lower levels in MNA tumours. We therefore assessed whether MYCN might directly control these genes. Consistent with MYCN repressing WMG-2 genes, depletion of MYCN by short-interfering RNA (siRNA) in SK-N-BE(2)-C cells led to a 2 – 8-fold induction of nine out of eleven WMG-2 genes tested, including *DCLK3*, *EPAS1*, *FAT4* and *NRP1* (Figure 7A, B). The effect of MYCN knockdown on WMG-1 and 3 genes was more variable, but *ETV6*, *BMP4* and *WNT11* were increased, and *NKD1*, *LIX1* and *NME2* transcripts were reduced as would be predicted by our clustering data.

**Figure 7.**
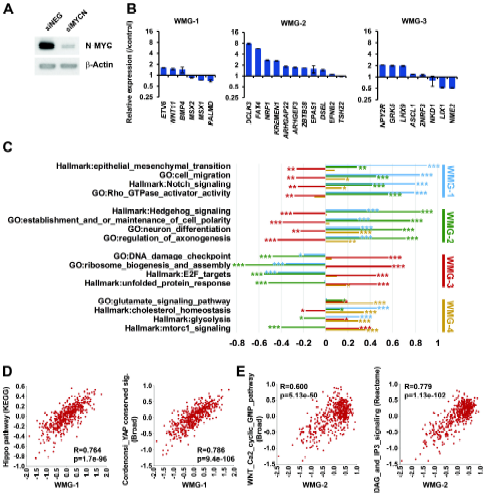
MYCN regulation of Wnt target genes. **(A)** Western blot of MYCN and β-actin, as loading control, 48 hours after *MYCN* knockdown in SK-N-BE(2)-C cells. **(B)** Changes in expression of Wnt target genes belonging to WMG-1, 2 and 3 following *MYCN* knockdown at 48 hours. **(C)** Correlation of Wnt metagenes with expression of curated gene sets at p<10^−10^ (***), p<10^−5^ (**) or p<0.01 (*). **(D)** Correlations with Broad Institute curated sets for WMG-1 **(E)** Correlations with Broad Institute curated sets for WMG-2.

We next analysed whether WMG-1, 2 and 3 are not simply sets of co-regulated genes, but whether they are also functionally linked. Interconnectivity analysis in the STRING protein-protein interaction database suggested that this was indeed the case, demonstrated by their significantly higher connections than expected by chance (Table 1). In contrast, genes in WMG-4 did not appeared to be functionally connected. The functional relevance of the Wnt metagene groups in neuroblastoma development was probed by enrichment analysis of Gene Ontology/Biological Process (GO/BP) categories. WMG-1 was found to be significantly enriched in 117 categories, broadly representing mesenchyme development, cellular migration, angiogenesis and regulation of proliferation. WMG-2 contained genes participating in sympathetic neuron projection and guidance, processes involved in neuronal differentiation, whereas WMG-3 and WMG-4 genes included several negative regulators of signalling pathways (*AMER1*, *ASCL1*, *AXIN2*, *DUSP4*, *ENC1*, *GRK5*, *NKD1*, *TNFRSF19*, *VLDLR*, and *ZNRF3*).

**Table 1.**
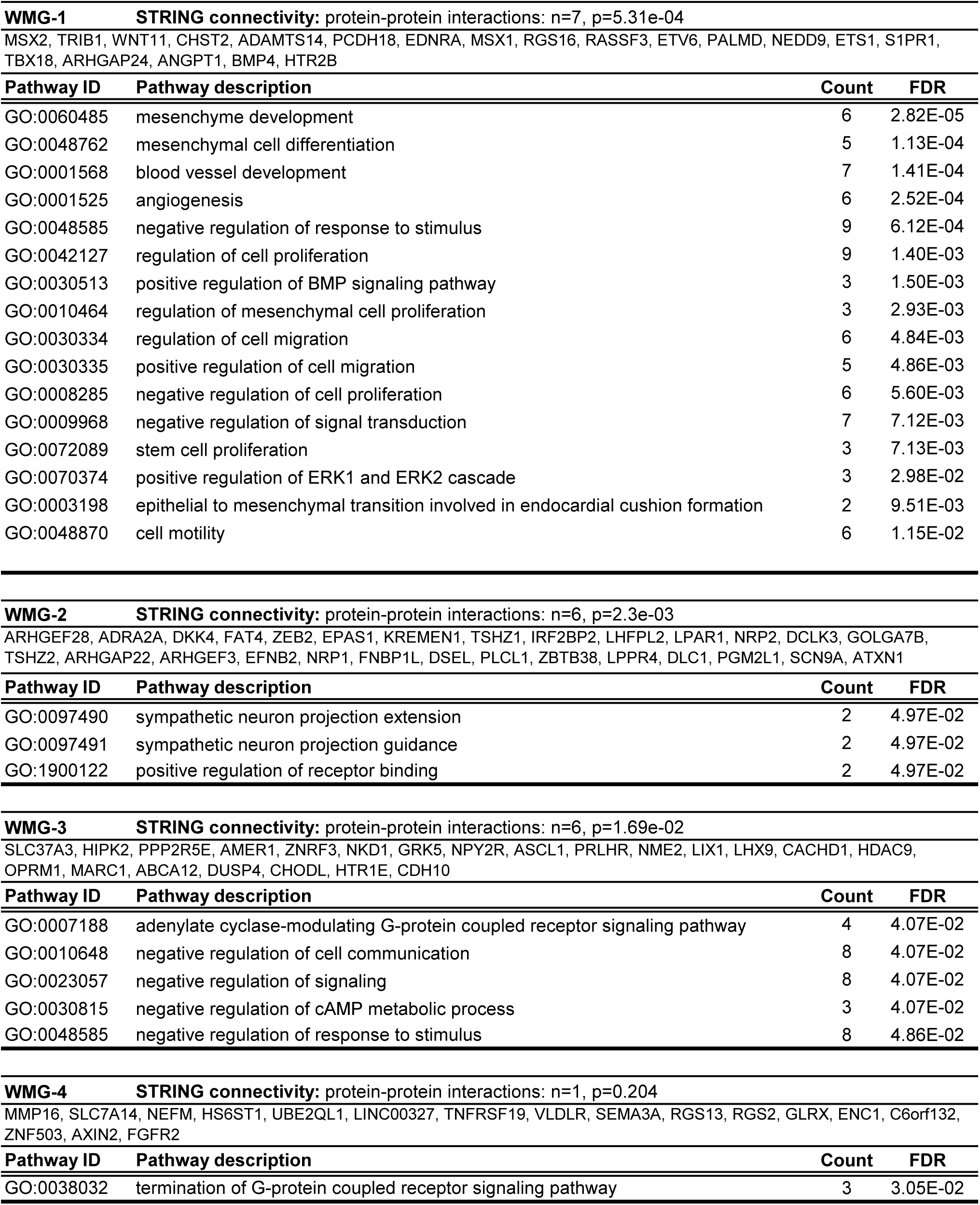
STRING and Gene Ontology analysis of Wnt metagenes.

We have already shown that Wnt3a/Rspo2 treatments could lead to decreases in other signalling pathways such as Akt and ERK signalling [26]. To further characterize signalling cross-talk and its possible biological significance, we searched for correlations between the metagenes and KEGG and Hallmark genesets (Figure 7C) and the Broad Institute curated gene sets (Figure 7D, E) within the neuroblastoma SEQC dataset. WMG-1 showed strong correlations with regulators of cell migration, EMT, Rho-GTPase activation and Notch signalling (Figure 7C), as well as Hippo/Yap signalling (Figure 7D). WMG-2 shared many of these correlations, but to a lesser extent. The highest correlations for WMG-2 was with gene sets driving neuronal differentiation, axonogenesis and maintaining cell polarity. Members of the Hedgehog pathway, an important driver of neural crest and neuroblastoma differentiation [52, 53], were also strongly associated with WMG-2 (Figure 7C). Broad genesets also revealed strong correlations of WMG-2 with alternative branches of the Wnt pathway such as the Wnt/Ca^2+^/cyclic GMP pathway and DAG/IP3 signalling (Figure 7E). WMG-3 showed strongest positive correlations for ribosome biogenesis, DNA damage check points, unfolded protein response and E2F target genes, which are all linked to high MYCN activity, and WMG-3 was generally anti-correlated with WMG-1 and WMG-2 gene sets. WMG-4 was associated with upregulation of metabolic pathways, including glycolysis, cholesterol homeostasis, glutamate pathway as well as mTORC1 signalling.

Together, these analyses suggest that expression of the Wnt metagene subsets is contingent on coregulatory axes. The most notable of these is MYCN suppression of WMG-2 genes, which likely represent genes critical for differentiation.

## Discussion

Wnt signalling is a key regulator of cell fate, capable of regulating stemness, differentiation and proliferation. The proliferative activity is often co-opted by cancer cells, acting via transcriptional transactivation of proto-oncogenes such as *MYC* and *CCND1*. In the case of neuroblastoma, Wnt signalling has been proposed to be an oncogenic driver of MYC over-expression in non-MNA tumours [24], although our previous work did not detect significant increases of c-MYC or MYCN protein after Wnt3a/Rspo2 treatment of neuroblastoma cell lines [26]. In this study, we therefore defined the Wnt-driven transcriptome in the MNA neuroblastoma cell line SK-N-BE(2)-C, and also examined the phenotypes accompanying Wnt signalling induced by recombinant Wnt3a and Rspo2 ligands. As well as revealing new Wnt targets likely to be involved in sympathetic nervous system development, and previously unrecognised signalling crosstalk in neuroblastoma, we demonstrate that Wnt signalling can prompt neuroblast differentiation, as well as proliferation of cell lines. Evaluation of the expression of our Wnt gene set in primary tumours reveals patterns of expression that are distinctive to different clinical cohorts and prognosis, indicating that, broadly, a suppression of Wnt signalling may be conducive to the development of high-risk neuroblastoma. This is further supported by immunohistochemical analysis of β-catenin staining, which is intense in ganglioneuromas and ganglioneuroblastomas and relatively weak in poorly differentiated neuroblastomas.

Of the 90 DEGs identified by our RNA-seq, 53 have previously been implicated as Wnt target genes by β-catenin and TCF4 binding to their promoters. We identified numerous transcription factors downstream of Wnt signalling, many of which have been shown to be involved in neuronal differentiation and to act upstream of Wnt signalling (*ASCL1* [42, 54], *MSX1* and *MSX2* [41], *LHX9* [55], and *TSHZ1* [56]). This, together with the other transcription factors (*EPAS1*, *ETS1*, *ETV6*, *TBX18*, *TSHZ2*, *ZEB2*, *ZBTB38*), as well as epigenetic regulators/non-coding RNAs (*HDAC9*, *KDM7A*, *LINC00327*, *TMEM108-AS1*, *VLDLR-AS1*) strongly supports Wnt signalling as a master regulator of neuroblast cell fate. In addition, crosstalk of Wnt signalling with several other signalling pathways can be inferred from our DEGs. These include MAPK (*DUSP4* [57], *HS6ST1* [58], *TRIB1* [59]), Hippo-Yap/Taz (*FAT4* [60], *LIX1* [61], semaphorin (*NRP1*, *NRP2*, *SEMA3A* [62]) and small G-protein/Rac/Rho signalling (*ARHGAP22*, *ARHGAP24*, *ARHGEF28*, *ARHGEF3, DLC1, FNBP1L* [63, 64]).

Consistent with the absence of obvious proto-oncogene induction within the DEGs, our examination of Wnt-induced phenotypes did not show a proliferative effect in SK-N-BE(2)-C cells. Instead, we observed clear neuritogenesis, concomitant with markers of EMT and neuronal differentiation. Here EMT likely reflects a transitional intermediate state with high plasticity that facilitates differentiation [65]. A similar neuronal differentiation phenotype was observed in SH-SY5Y cells lines, but in the case of SK-N-AS cells, we observed Wnt-induced proliferation, as recently shown [23]. At the molecular level, the different responses of these cell lines were accurately reflected by changes in known drivers of differentiation (BMP4, EPAS1, NGFR) and proliferation (CCND1, E2F1 and phospho-RB). To our knowledge this is the first demonstration of Wnt ligands inducing phenotypic differentiation of neuroblastoma cells, in particular MNA neuroblastoma cells. Consistent with our results, a very recent study suggested that the tankyrase inhibitor XAV939 reduces markers of differentiation in SH-SY5Y cells, although in contrast to our work, that study observed no effects on morphology or proliferation [66]. In another report using chemical Wnt agonists and inhibitors, Wnt inhibition was found to block proliferation and promote neuroblastoma differentiation, and Wnt induction led to cell death [25]. The reasons for these discrepancies are unclear, but may be attributable to the cell-line models used, different dosing regimens and additional non-specific effects of chemical Wnt modulators. Consistent with this previous report, however, we did observe MYCN and Wnt interaction, although the genes we show to be MYCN repressed have not previously been demonstrated to be MYCN-regulated.

Whilst the differentiating effects of Wnt ligands may be significant for potential differentiation therapies, the proliferative effect seen with SK-N-AS cells underlines the need for further delineation of the genotypes and transcriptional/signalling contexts determining Wnt responses. SK-N-AS cells are also known to be resistant to ATRA-mediated differentiation [67]. As well as having mutant *NRAS*, SK-N-AS cells have been shown to be dependent on Taz for proliferation [68]; as both Ras and Hippo-Yap/Taz can associate with Wnt signalling, we examined Ras activity and Taz protein levels in SK-N-AS cells after Wnt induction, but did not observe any differences (data not shown). Two recent studies have shown that SK-N-AS cells have a transcriptional context that is quite different from most neuroblastoma cell lines [4, 5], with the first of these dividing neuroblastoma cells into mesenchymal (MES) or adrenergic (ADRN) types. Our analysis of another MES line, GI-MEN, did not however support the notion that induction of proliferation by Wnt is solely attributable to a MES identity. We have previously shown that SK-N-AS cells are also quite different from SK-N-BE(2)-C and SH-SY5Y cells in terms of their genome-wide DNA methylation pattern [69], and the epigenetic identity of SK-N-AS has been shown to influence their differentiation response [28, 70]. Thus further analysis at the epigenetic and proteomic level with a broad range of neuroblastoma cell lines will be necessary to precisely identify the determinants of the phenotypic responses to Wnt signalling by neuroblastoma cells.

To date, several studies have proposed that elevated expression of Wnt/β-catenin target genes can be used to derive a prognostic signature indicative of poor prognosis in neuroblastoma [24, 25, 49]. However, analysis of primary tumour datasets shows that expression of our *bona fide* Wnt gene set, which has essentially no overlap with gene sets used in the other studies, reveals a much more complex picture, with expression patterns varying in four distinct patient cohorts. Low expression of the metagene WMG-2 is a strong indicator of poor survival and conversely elevated expression of WMG-3 is associated with poor prognosis, and is particularly high in the predominantly MNA cohort. Importantly, both WMG-2 and WMG-3 identify patients at greater risk despite being classified as low risk patients, with WMG-2 showing consistent and robust statistical significance across neuroblastoma expression datasets. Remarkably, although Asgharzadeh *et al*. suggested that high expression of a Wnt signature (based on existing non-neuroblastoma Wnt datasets) is associated with poor prognosis, we found that WMG-2 shows quite the opposite trend, and in fact has a striking positive correlation with the powerful Asgharzadeh signature, whose low expression correlates with poor prognosis in non-MNA neuroblastoma [49]. This further underlines the value of our empirically determined neuroblastoma Wnt target gene set. Whilst evaluation of the power and utility of Wnt signatures for prognosis is beyond the scope of this study, further determination of Wnt target genes in neuroblastoma models may yield powerful prognostic signatures.

The inference from our work that Wnt signalling may have a pro-differentiation and tumour suppressive influence is supported by the recent identification of Wnt pathway mutations in neuroblastoma, which include probable loss of function mutations of *LEF1* and *VANGL1* [20]. Comparison of our 90 DEGs with mutations identified in this study further revealed that 9 of our Wnt targets (10%) also exhibited mutations in neuroblastoma; these included *ADAMTS14*, *AXIN2*, *CACHD1*, *DUSP4*, *IRF2BP2*, *NPY2R*, *NRP2*, *OPRM1*, and *ZNRF3*. In addition, *LGR5* was also mutated in one sample, adding to the previously reported *LGR5* mutations [13]. Although the pathological consequences of these mutations remains to be established, these studies, together with ours, suggest that impairment of Wnt signalling can contribute to neuroblastoma tumourigenesis.

Numerous regulatory pathways that likely interact/crosstalk with Wnt/β-catenin signalling emerged through correlation analyses of our Wnt metagenes. WMG-2 genes, which appear to be down-regulated in MNA tumours, could be derepressed after MYCN knockdown in SK-N-BE(2)-C cells, suggesting they are direct targets of MYCN. An inverse correlation of *MYCN* and *EPAS1* was suggested recently [28], and consistent with this *EPAS1* increased after MYCN knockdown (Figure 5). The largest change observed was doublecortin-like kinase 3 (*DCLK3*), and although the functions of DCLK3 are not characterised, we note that its paralogue DCLK1 has been shown to be vital in neuronal migration and axon outgrowth [71]. This is in keeping with the overall trend for WMG-2 to correlate with neuron differentiation and axonogenesis, and its associated pathways, including Hedgehog [53] and Rho-GTPase activation [64]. Notably, genes encoding regulators of Rac/Rho signalling, were reported to be mutated in a whole-genome sequencing analysis of 87 neuroblastomas, including one of our WMG-2 genes *DLC1* (Deleted In Liver Cancer 1/ARHGAP7) [72]. Together this suggests that loss of Wnt stimulation of Rac/Rho signalling (via MYCN-mediated repression) adds to the mutational disruption of neuritogenesis.

Another key factor that may underlie the expression patterns we observe in primary tumours is the tumour microenvironment. For example, Schwannian stroma is associated with better prognosis of neuroblastoma [50] and can induce differentiation of neuroblasts [73]. Our immunohistochemistry indicates that well-differentiated/differentiating tumours have high levels of β-catenin in the Schwannian stroma, and it is therefore tempting to speculate that Wnt signalling may induce paracrine differentiating factors such as BMP4 and Wnt11 to prompt differentiation of neuroblast cells by Schwann cells. Our future work will aim to investigate this possibility and further dissect the complex signalling crosstalk in neuroblastoma using co-culture organoid systems and *in vivo* models.

## Conclusion

Our study reveals that Wnt signalling can promote differentiation of neuroblastoma via orchestrating a myriad of transcriptional and signalling nodes, and suggest that Schwannian stroma may be a source of differentiation signals. Therefore highly specific Wnt small molecule agonists may have utility in differentiation therapies for neuroblastoma in the future. However, our data also underlines the problems of cellular heterogeneity in neuroblastoma, with SK-N-AS cells proliferating rather than differentiating. The challenge of defining critical regulatory circuits controlling differentiation in the context of the genetic and signalling heterogeneity of neuroblastoma is immense, but our study emphasises that Wnt signalling is one of the key players.

## Acknowledgements

We thank Keith Brown for helpful discussions, Gabriella Cunha Vieira for assistance with sample preparation, and Melanie Panagi for assistance with wound/scratch assays. This work was assisted using the computational facilities of the Advanced Computing Research Centre, University of Bristol. We would like to thank Dr. Jan Kosters and team (Academic Medical Center, Amsterdam) for R2 tuition, and Children with Cancer UK, Neuroblastoma UK and Smile with Siddy, the Childrens Cancer and Leukaemia Group (CCLG), the Biotechnology and Biological Sciences Research Council (BB/P008232/1) and Cancer Research UK (A12743/A21046) (to K.M.) for funding this study.

